# Central hormone-sensitive lipase is located at synapses and is necessary for normal memory performance in mice

**DOI:** 10.1101/2021.09.03.458841

**Authors:** Cecilia Skoug, Cecilia Holm, João M.N. Duarte

## Abstract

Hormone-sensitive lipase (HSL) is mainly present in the adipose tissue where it hydrolyses diacylglycerol. Although brain expression of HSL has been reported, its presence in different cellular compartments is uncertain, and its role in regulating brain lipid metabolism remains hitherto unexplored. We propose that HSL has a role in regulating the availability of bioactive lipids necessary for adequate neuronal function. Therefore, we tested the hypothesis that dampening HSL activity leads to brain dysfunction. We found HSL protein and activity throughout all the mouse brain, localised in neurons and especially enriched in synapses. HSL null mice were then analysed using a battery of behavioural tests. Relative to wild-type littermates, HSL null mice showed impaired short- and long-term memory, but preserved exploratory behaviours. Molecular analysis of the cortex and hippocampus showed increased expression of genes involved in glucose utilization in the hippocampus but not cortex of HSL null mice compared to controls. Lipidomics analyses indicated an impact of HSL deletion on the profile of bioactive lipids, including endocannabinoids and eicosanoids that are known to modulate neuronal activity, cerebral blood blow and inflammation processes. Accordingly, mild increases in expression of pro-inflammatory cytokines suggest low grade inflammation in HSL null mice compared to littermates. We conclude that HSL has a homeostatic role in maintaining pools of lipids that are needed for brain function. It remains to be tested, however, whether the recruitment of HSL for the synthesis of these lipids occurs during increased neuronal activity, or whether HSL participates in neuroinflammatory responses.

## Introduction

Breakdown of triacylglycerols (TAG) in adipose tissue is under hormonal regulation and is mediated by adipose triglyceride lipase (ATGL) that hydrolyses a fatty acid from TAG generating DAG, then the hormone-sensitive lipase (HSL) hydrolyses diacylglycerols (DAG), and finally monoacylglycerol lipase (MAGL) degrades monoacylglycerols into a fatty acid and glycerol. HSL is abundant in the adipose tissues where it mainly hydrolyses DAG, and plays a key role in lipolysis regulation since it is stimulated by catecholamines and inhibited by insulin (Haemmerle *et al*., 2002).

The brain, like most organs, utilises fatty acids as oxidative substrate (Kuge *et al*., 1995; Ebert *et al*., 2003). The brain expresses ATGL (Etschmaier *et al*., 2011) and HSL (Haemmerle *et al*., 2002), which have been suggested to control the release of free fatty acids to be used as energy substrate. Although brain cells can oxidise alternative substrates under a variety of physiological or pathological conditions, glucose is the obligatory fuel for adequate brain function (Sonnay *et al*., 2017; Dienel, 2019). Therefore, HSL is unlikely to be necessary for regulating fatty acid availability from acylglycerols for local energy production.

MAGL activity is ubiquitous in the brain and has a pivotal role in the endocannabinoid system: it terminates the action of the main endocannabinoid 2-arachidonoylglycerol (2-AG) through hydrolysis into glycerol and arachidonic acid (Blankman *et al*. 2007). This is an example in which the lipase is not fulfilling energetic needs of brain cells, but contributing to functional control of neuronal activity. Endocannabinoids control brain metabolism and synaptic function (Metna-Laurent & Marsicano, 2015). The endocannabinoid 2-AG is synthetized by diacylglycerol lipase (DAGL) α and β at postsynaptic terminals following stimulation by glutamate, and is then released to presynaptic terminals where it stimulates cannabinoid 1 receptors, thus dampening glutamatergic activity at synapses (Iannotti *et al*., 2016). Moreover, cannabinoid 1 receptors directly inhibit mitochondrial metabolism in neurons and astrocytes (Duarte, Ferreira *et al*., 2012), which may contribute to 2-AG-dependent synaptic control. Interestingly, while having negligible 2-AG hydrolysis activity (Blankman *et al*., 2007), HSL could rather fulfil a role in hormone-regulated 2-AG production, thus integrating signalling from catecholamines and cannabinoids at neuronal synapses.

The main endocannabinoids 2-AG and *N*-arachidonoylethanolamine (or anandamide) are quickly hydrolysed by MAGL (Dinh *et al*, 2002) and fatty acid amide hydrolase (FAAH) (Cravatt *et al*, 1996), respectively. These reactions produce arachidonic acid that, in turn, is enzymatically oxidized to produce eicosanoids that are pro-inflammatory mediators (Donvito *et al*., 2018) and modulators of cerebral blood flow (Atwell *et al*., 2010; Sonnay *et al*., 2017). Thus, DAG hydrolysis by HSL could also contribute to regulate metabolic and signalling pools of arachidonic acid and other bioactive lipids.

Given the ability of HSL to be controlled by multiple cellular signals (Kraemer & Shen, 2002), we propose that its activity in the brain is poised to regulate lipid metabolism in cellular compartments where lipolysis is stimulated on demand for release of precursor fatty acids from esterified forms, which are then used for production of bioactive lipids. In this study we aimed at (i) mapping the relative distribution of HSL in the rodent brain, and (ii) investigating the impact of HSL deletion on brain function, namely memory performance, and exploring underlying mechanisms.

## Methods

### Animal models

All procedures on animals were approved by the Malmö-Lund Committee for Animal Experiment Ethics and conducted according to EU Directive 2010/63/EU, and are reported following the ARRIVE guidelines (Animal Research: Reporting In Vivo Experiments, NC3Rs initiative, UK). Mice were housed on a 12-hour light-dark cycle with lights on at 7:00, room temperature of 21-23 °C and humidity at 55-60%, and had access to regular chow and water *ad libitum*. Male C57BL/6J mice were obtained from Taconic (Ry, Denmark) at eight weeks of age and allowed to acclimatize to the animal facility for at least 2 weeks before experiments. HSL-null (HSL-/-) mice were generated by targeted disruption of the HSL gene in 129SV-derived embryonic stem cells as described elsewhere (Mulder *et al*., 2003) and back-crossed to a C57BL/6J background for nine generations. HSL-/- mice and wildtype +/+ littermates of either gender were studied at 19-22 months of age.

### Neuronal cell cultures

Primary embryonic neurons were prepared from the cortices and hippocampi of embryonic day 15-17 WT mouse embryos, as detailed by Martinsson *et al*. (2019). Briefly, neurons were dissociated through trypsinization and subsequent trituration in Dulbecco’s modified Eagle medium (DMEM, 30243.01#, Cytiva) supplemented with 10% fetal bovine serum (#10100-147, Gibco, Australia), 1% penicillin-streptomycin (#15140122, Thermo fischer) and then placed onto poly-D-lysine coated coverslips. After 3-5 hours, medium was switched to Neurobasal medium supplemented with glutamine (#25030081, Thermo fischer), B27 (#A3582801, Thermo fischer), pencilin-streptomycin. Cells were grown *in vitro* for 19-20 days and then were fixed for 15 minutes in 4% paraformaldehyde and 4% sucrose at room temperature, and stored in phosphate-buffered saline (PBS; in mmol/L: 137 NaCl, 2.7 KCl, 1.5 KH_2_PO_4_, 8.1 Na_2_HPO_4_, pH 7.4) at 4 °C until usage.

### Behavior testing

Mice were allowed to acclimatize to the testing room for 1 hour before each experiment, and tests were performed from 9:00 to 18:00. Barnes maze tests were conducted under bright light over the platform. Otherwise, room light was adjusted to an illuminance of 15 lx in the test apparatus. Experiments were recorded by an infrared camera into the AnyMaze software (6.0.1, Stoelting).

A circular Barnes Maze with diameter of 92 cm and 20 wholes placed at a height of 90 cm was used to test learning and memory (Attar *et al*., 2003). The target hole had a removable dark escape box under the maze, and four proximal visual cues were place at 20 cm from the platform. Experiments consisted of habituation, 8-day acquisition (training) and memory retention trial. For the habituation, mice were placed in the escape box during 60 s, and then released in the center of the apparatus, and allow it to explore until re-entering the escape box, or until 5 minutes elapsed. The first acquisition trial was conducted two hours after the habituation, and was used to probe short-term memory. Acquisition trials took place in 8 consecutive days, at the same time of the day, in which mice were released in the center of the maze with head pointing in random direction, and allowed to explore the maze for 5 minutes. The test ended when mice escaped into the target-hole box. Whenever mice did not find the escape box within 5 minutes, they were gently guided into it. The memory retention trial took place 48 hours after the last acquisition session, under identical conditions but in the absence of any escape box. To eliminate olfactory cues, the surface and the escape box were cleaned with 70% (v/v) ethanol in-between each trial. Measured parameters were path and latency to reach or enter the target hole, and time spent in each quadrant of the maze during retention trial (Harrisson *et al*., 2006).

Spontaneous alternations were observed in a Y-maze as surrogate of working memory performance (Duarte, Agostinho *et al*, 2012). The Y-maze arms were 30 cm x 15 cm x 5 cm (length x height x wide), and converged to the center in 120° angles. Mice were placed an arm of the maze and allowed to freely explore for 5 minutes. A complete spontaneous alternation was defined as a successive entrance to each different arm, and expressed relative to the total possible alternations in the respective test. The total number of entries was analysed to access locomotor activity and exploratory behaviour.

Open-field exploration was recorded for 5 minutes in a cubic arena with length of 50 cm by an infrared camera. Arena exploration was analysed for total walk distance, number of crossings between arena quadrants and number of rearing events, as well as exploration of the arena center at 6 cm from the walls.

The Elevated Plus Maze was used to assess anxiety. Each maze arm was 35 cm x 5 cm, and closed arms had 15 cm walls, at a 60 cm height from the floor. The mouse was placed in the maze center facing an open arm, and was allowed to freely explore the maze for 5 minutes. Number of entries and time spent in each arm were analysed (Walf *et al*., 2007).

### Tissue sampling

Mice were anaesthetized with isoflurane and quickly decapitated, and brains were dissected, frozen in N_2_ (l) and stored at -80 °C until further experiments. For immunofluorescence experiments, mice under isoflurane anaesthesia were sacrificed by cardiac perfusion with cold PBS and then cold phosphate-buffered formaldehyde (Histolab, Askim, Sweden), and brains were processed as detailed previously (Mohr *et al*., 2020).

### Plasma glucose and insulin

Commercially available ELISA kit were used to determine plasma concentration of insulin (#10-1247-10, Mercodia, Sweden). Glucose plasma concentration was measured with glucose oxidase method using 2’2-azino-bis (3-ethylbenzothialozine-6-sulfonate) (ABTS) (#10102946001, Roche, Germany) as substrate. The absorbance was measured at 420nm on a microtiter plate reader (Floustar/Polystar Galaxy, BMG Labtech, Germany)

### HSL activity

Tissues were mechanically homogenized in 0.25 mol/L sucrose, 1 mmol/L ethylenediaminetetraacetic acid (EDTA), 1 mmol/L dithiothreitol and protease inhibitors (20 μg/mL leupeptin, 2 μg/mL antipain, 1 μg/mL pepstatin), pH 7.0, using a glass-glass homogenizer. To remove fat from the tissues, it was centrifuged at 110 000×g for 1 hour at 4°C (Optima TLX Ultracentrifuge, Beckman) and the fat-free infranatant was used for HSL activity measurements. Measurement of HSL activity was performed using 1 (3)-oleoyl-2–0-oleylglycerol, a diacylglycerol analogue, as substrate, as described in detail by Holm and Osterlund (1999). It should be emphasized that the diacylglycerol analogue used as substrate, does not generate substrate for MAGL due to the ether bond in position 2. To estimate the fraction of activity accounted for by HSL, samples were pre-incubated with either an activity-neutralizing hen anti-rat HSL serum (prepared in-house) or pre-immune serum for 20 minutes at 37°C prior to the assay. Total protein content of the samples was measured with the bicinchoninic acid assay (kit from Pierce, Thermofisher Scientific, Göteborg, Sweden). Activity was expressed as mU/mg protein, where 1 U corresponds to the release of 1 μmol of fatty acids per min at 37°C.

### Total protein extracts

Tissue samples were homogenized with a sonicator probe in lysis buffer (in mmol/L: 150 NaCl, 1 EDTA, 50 tris(hydroxymethyl)aminomethane (Tris)-HCl, 1% (w/v) sodium dodecylsulfate (SDS), pH 8.0) containing protease inhibitors (#11697498001, Roche, Switzerland) and phosphatase inhibitors (#4906837001, Roche, Switzerland). The homogenate was maintained in constant agitation for 2 hours at 4 °C. After centrifugation at 3,000 *g* for 10 minutes at 4 °C to remove major debris, the supernatant was saved. Protein concentration was determined as described above.

### Preparation of synaptosomes and synaptic fractions

Synaptosomal fractionation was modified from Morato *et al*., 2017. Briefly, mouse cortex was homogenized in 1 mL of isolation buffer (in mmol/L: 320 sucrose, 0.1 CaCl_2_, 0.1 MgCl_2_, pH 7.4) at 4°C in a 5-mL Potter-Elvehjem glass/teflon homogenizer (10 strokes at 700–900 rpm). The resulting homogenate was mixed with 6 mL sucrose (2 mol/L) and 2.5 mL CaCl_2_ (0.1 mmol/L) in an ultra-clear centrifuge tube (#344059, Beckman Coulter, USA). Then, 2.5 mL of sucrose (1 mol/L) containing 0.1 mM CaCl_2_ were carefully added on top to form a discontinuous sucrose gradient. All centrifugations were performed in an Optima XL-100K Ultracentrifuge (Beckman Coulter) with SW41Ti swinging bucket rotor (Beckman Coulter). After centrifugation for 3 hours at 100,000 *g*, 4°C, the synaptosomes were collected from the interphase between 1.25 and 1 mol/L sucrose and diluted 10 times in isolation buffer, centrifuged for 30 minutes at 15,000 *g*, 4°C, and the resulting synaptosomal pellet was re-suspended in 1 mL of isolation buffer.

For fractioning synaptosomes, part of each sample was diluted 1:5 in 0.1 mmol/L CaCl_2_, and an equal volume of solubilization buffer (2% Triton X-100, 40 mmol/L Tris, pH 6.0) was added to the suspension. The suspension was incubated for 30min on ice with constant agitation and the insoluble material (synaptic junctions) was pelleted by centrifugation for 30 minutes at 40,000 *g*, 4°C. The supernatant (extra-synaptic fraction) was concentrated using an Amicon Ultra 15 10K (#UFC901008, Merck Millipore, Ireland) and protein was precipitated with six volumes of acetone at -20 °C and recovered by centrifugation for 30 minutes at 18,000 *g*, -15 °C. The pellet containing synaptic junctions was washed in solubilization buffer at pH 6.0, and then re-suspended in 10 volumes of a second solubilization buffer (1% Triton X-100 and 20 mmol/L Tris, pH 8.0). After incubation under agitation for 30 minutes on ice, the mixture was centrifuged and the supernatant (pre-synaptic fraction) was processed as described for the extra-synaptic fraction, whereas the insoluble pellet corresponds to the post-synaptic fraction All synaptic fractions were resuspended in 5% SDS with protease inhibitors.

### Immunoblotting

Western blotting was carried out as previously reported (Lizarbe et al., 2019). Briefly, samples were heated for 5 minutes at 95°C in sample buffer (#NP0007, Invitrogen, USA), and then separated on 4-12% Bis-Tris mini gels (#NP0336, Invitrogen, USA), followed by transfer onto nitrocellulose membranes, pore size 0,45 μm (#GE10600002, GE Healthcare, Germany). The membranes were blocked for 60-120 minutes in 5% milk or bovine serum albumin in Tris-buffered saline (in mmol/L: 20 Tris, 150 NaCl, pH 7.6) containing 1% Tween 20, and incubated with primary and secondary antibodies (table 3) diluted in this blocking solution. Immunoblots were developed with a chemiluminescence kit (#34580, Thermo Scientific, USA) using the Chemidoc XRS+ interfaced to Image Lab 5.2.1 for image analysis (Biorad, Stockholm, Sweden).

**Table 1.**
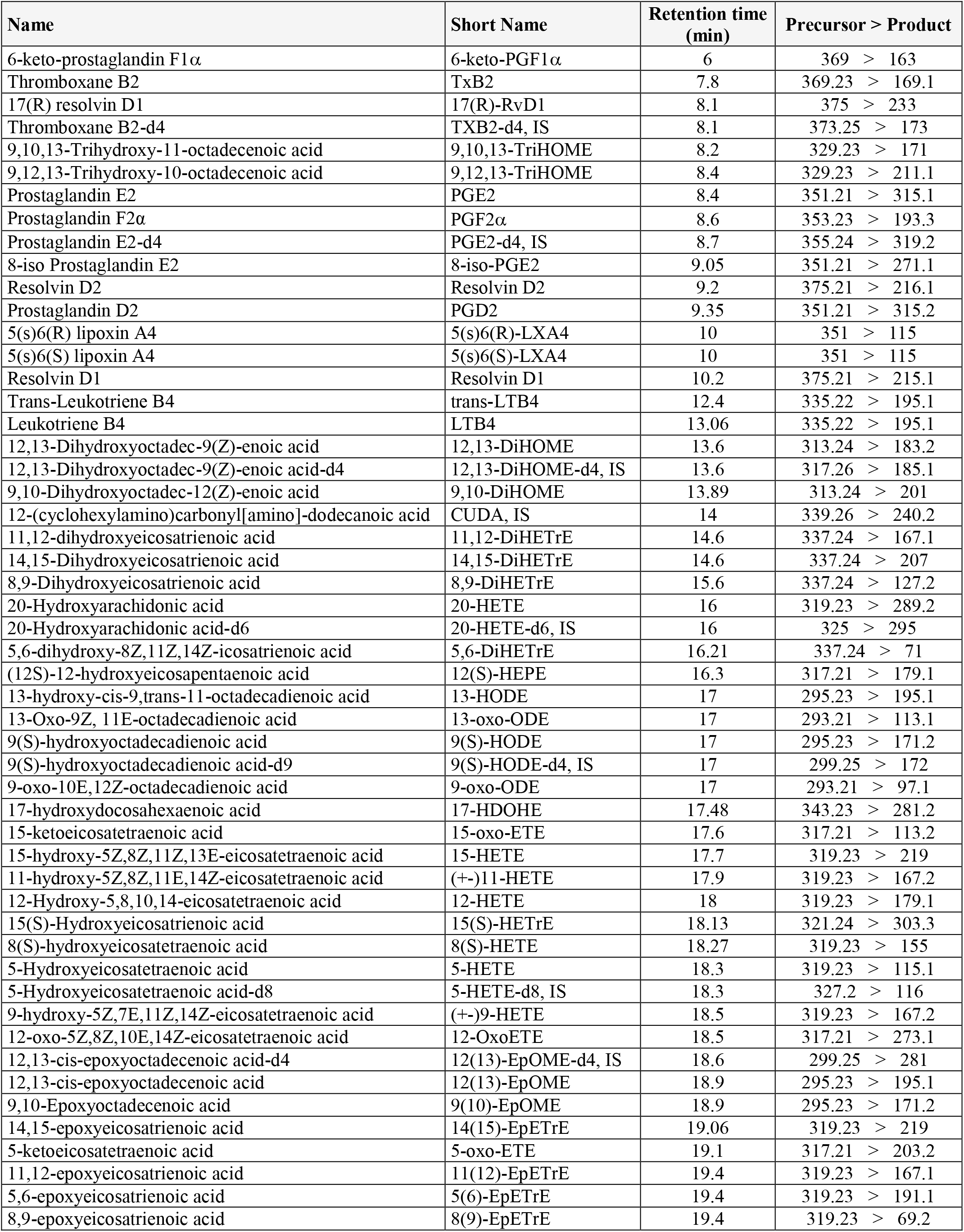
Oxylipins in the lipidomics analysis, their retention times and multiple reaction monitoring transitions. IS, internal standards.

**Table 2.** Endocannabinoids in the lipidomics analysis, their retention times and multiple reaction monitoring transitions

| Name | Short Name | Retention time (min) | Precursor > Product |
| --- | --- | --- | --- |
| Prostaglandin F2α ethanolamide | PGF2α EA | 3.19 | 380 > 62 |
| Prostaglandin F2α ethanolamide-d4 | PGF2α EA-d4, IS | 3.19 | 384 > 62 |
| Prostaglandin E2 ethanolamide | PGE2 EA | 3.24 | 378 > 62 |
| 12-(cyclohexylamino)carbonyl[amino]-dodecanoic acid | CUDA, IS | 4.32 | 341 > 216 |
| Eicosapentaenoyl ethanolamide | EPEA | 5.66 | 346 > 62 |
| Palmitoleoyl ethanolamide | POEA | 6.27 | 298 > 62 |
| Docosahexaenoyl ethanolamide | DHEA | 6.31 | 372 > 62 |
| Arachidonoyl ethanolamide | AEA | 6.58 | 348 > 62 |
| Arachidonoyl ethanolamide-d8 | AEA-d8 | 6.58 | 356 > 63 |
| N-Arachidonoyl glycine | NAGly | 6.66 | 362 > 287 |
| Linoleoyl ethanolamide | LEA | 6.72 | 324 > 62 |
| 2-Arachidonoyl glycerol-d8 | 2-AG-d8, IS | 7.22 | 387 > 294 |
| 2-Arachidonoyl glycerol | 2-AG | 7.3 | 379 > 287 |
| 2-Linoleoyl glycerol | 2-LG | 7.63 | 355 > 263 |
| Palmitoyl ethanolamide-d4 | PEA-d4, IS | 7.99 | 304 > 62 |
| Palmitoyl ethanolamide | PEA | 8.04 | 300 > 62 |
| Docosatetraenoyl ethanolamide | DEA | 8.21 | 376 > 62 |
| Oleoyl ethanolamide-d4 | OEA-d4, IS | 8.48 | 330 > 66 |
| N-Oleoyl ethanolamine | OEA | 8.52 | 326 > 62 |
| Stearoyl ethanolamide | SEA | 11.85 | 328 > 62 |
| Stearoyl ethanolamide-d3 | SEA-d3, IS | 11.85 | 331 > 62 |

**Table 3.** Key resources table.

| REAGENT or RESOURCE | SOURCE | IDENTIFIER (RRID/Cat#) |
| --- | --- | --- |
| **Antibodies** |  |  |
| Mouse monoclonal anti-β-actin | Sigma-Aldrich | AB_262011 |
| Rabbit monoclonal anti-GFAP | Abcam | AB_880202 |
| Affinity purified chicken anti-HSL | In-house | AB_2892188 |
| Rabbit polyclonal anti-Iba1 | FUJIFILM Wako | AB_839506 |
| Rabbit anti-Syntaxin 1 | Sigma-Aldrich | AB_261426 |
| Rabbit monoclonal anti -Syntaxin 4 | Abcam | AB_2891056 |
| Rabbit monoclonal anti-SNAP25 | Abcam | AB_10887757 |
| Rabbit monoclonal anti-Synaptophysin | Abcam | AB_2286949 |
| Hen anti-rat HSL serum | In-house | AB_2892189 |
| Affinity purified rabbit anti-HSL | In-house | AB_2892190 |
| Rabbit monoclonal anti-MAP2- alexa 488 tagged | Abcam | AB_2891057 |
| Chicken polyclonal anti-MAP2 | Abcam | AB_2138153 |
| Mouse monoclonal anti-NeuN | Abcam | AB_10711040 |
| Mouse monoclonal anti-PSD95 | Millipore | AB_2868387 |
| **Chemicals, Peptides, and Recombinant Proteins** |  |  |
| protease inhibitors | Roche | #11697498001 |
| phosphatase inhibitors | Roche | #4906837001 |
| sample buffer | Invitrogen | #NP0007 |
| Bis-Tris mini gels | Invitrogen | #NP0336 |
| goat serum | Gibco | #16210-064 |
| PerfeCTa SYBRgreen SuperMix | Quantabio | #95054 |
| qScript cDNA SuperMix | Quantabio | #95048 |
| ProLong Glass Antifade | Invitrogen | #P36980 |
| 2'2'-azino-bis (3-ethylbenzothiazoline-6-sulfonate) (ABTS) | Roche | #10102946001 |
| **Critical Commercial Assays** |  |  |
| Insulin ELISA kit | Mercodia | AB_2783837 |
| Chemiluminescence kit | Thermofisher | #34580 |
| Bicinchoninic acid kit | Thermofischer | #23227 |
| **Experimental Models: Organisms/Strains** |  |  |
| HSL null mice | Mulder et al., 2003 | N/A |
| **Oligonucleotides** |  |  |
| Primers (see table 4) | This paper | N/A |
| **Software and Algorithms** |  |  |
| AnyMaze 6.0.1 | Stoelting | SCR_014289 |
| Image Lab 5.2.1 | Biorad | SCR_014210 |
| ImageJ 1.53j | NIH | SCR_003070 |
| NIS-elements 5.20.01 | Nikon | SCR_014329 |
| Primer-BLAST | NIH | SCR_003095 |
| MassHunter Quantitative Analysis QQQ | Agilent | SCR_015040 |
| Prism 9.0.2 | GraphPad | SCR_002798 |
| MATLAB 2019a | MathWorks | SCR_001622 |

### Immunofluorescence confocal microscopy

Primary neurons on cover glasses or 20-µm cryostat-sectioned coronal brain slices were incubated for 1 hour at room temperature with blocking buffer [PBS containing 5% (v/v) goat serum (#16210-064, Gibco, New Zealand), 1% (w/v) bovine serum albumin and 0.3% (v/v) Triton X-100], followed by 2-hour incubation with primary antibodies (table 3). After washing in PBS, the samples were incubated with AlexaFluor-conjugated secondary antibodies, washed again, mounted for microscopy with ProLong Glass Antifade (#P36980, Invitrogen, USA), and examined under a Nikon A1RHD confocal microcope with a CFI Apochromat TIRF 100x Oil, NA 1.49 or CFI Plan Apochromat Lambda 20x NA 0.75 (Nikon Instruments, Tokyo, Japan). Images were acquired with NIS-elements (Laboratory Imaging, Nikon), and then processed in ImageJ (NIH, Bethesda, MD, USA).

### Real-time polymerase chain reaction (RT-PCR)

RNA was isolated from cortex and hippocampus from one hemisphere using Trizol (#15596026, Invitrogen, USA), and then 1 μg of total RNA was reverse transcribed with random hexamer primers using the qScript cDNA SuperMix (#95048, Quantabio, England), according to the manufacters’ instructions. The resulting cDNA was used as template for quantitative RT-PCR in triplicates using PerfeCTa SYBRgreen SuperMix (#95054, Quantabio, England) and the primers in table 4. Cycling and detection were carried out using Quantstudio 5 Real-time PCR system (40 cycles of 5 s at 95 °C and 30 s at 60 °C). Primers were optimized prior usage by assessing the optimal annealing temperature. Specificity was monitored using melt-curve validation and confirmation of amplified product by agarose gel electrophoresis. The dynamic range of each PCR assay was determined by constructing a standard curve using serial dilution of cDNA representative of each sample. Samples and standards were run in triplicate. All data were normalized to the expression of 60S ribosomal protein L14 and analyzed with the comparative threshold cycle method (ΔΔCT).

**Table 4.**
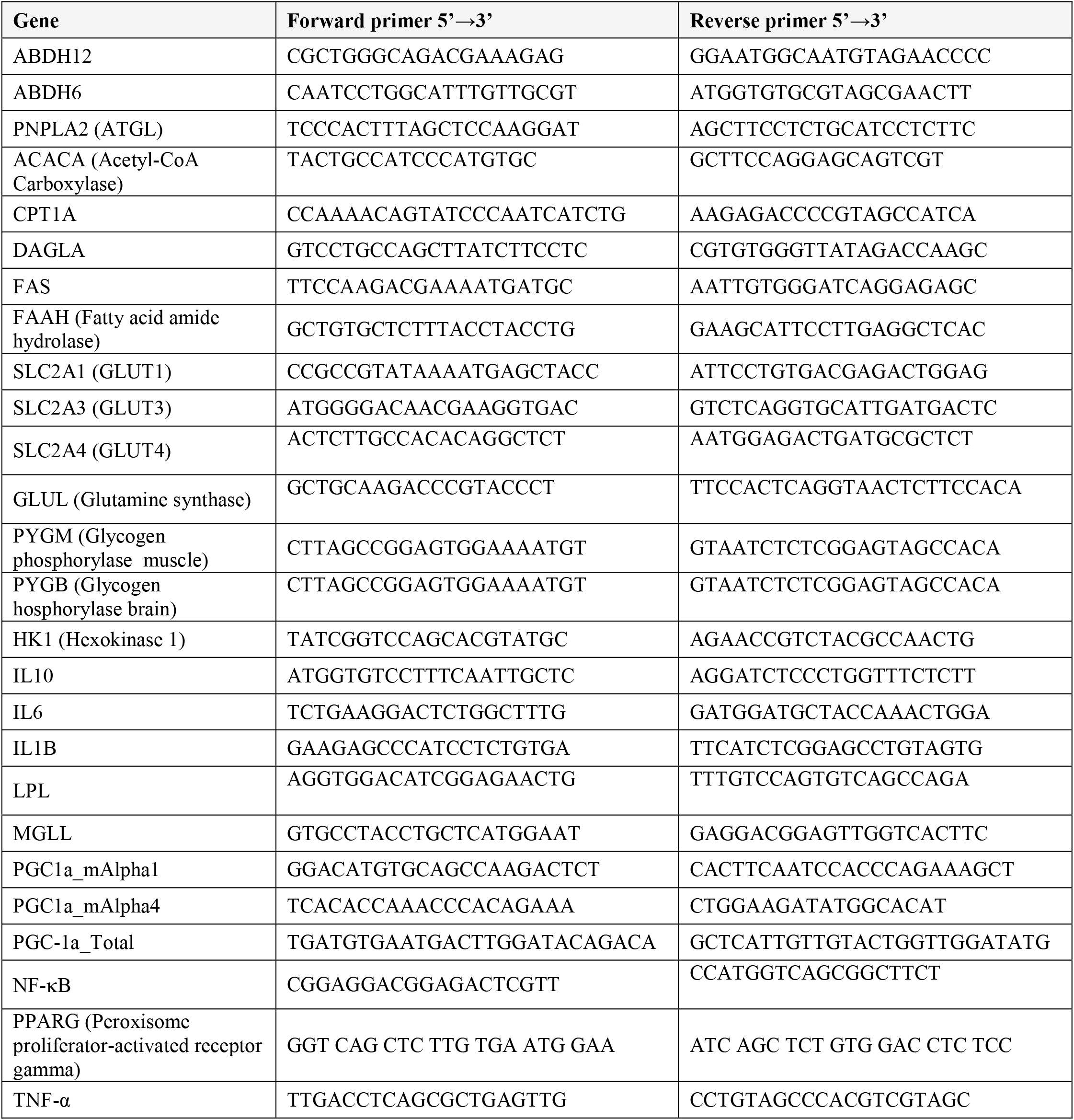
Primers used for RT-PCR.

### Lipidomics

Analysis of oxylipins (table 1) and endocannabinoids (table 2) in tissue by UHPLC-QqQMSMS was carried out as detailed before (Gouveia-Figueira and Nording, 2015). Briefly, 500 µL methanol was added to ∼20 mg of sample material. The sample was shaken with two tungsten beads at 30 Hz for 3 min in a mixer mill (MM 400, Retsch) whereupon the beads were removed. Samples were centrifuged at 2,125 *g* for 10 minutes at 4 °C and 450 µL of the supernatant was diluted in 8.5 mL of 0.1% acetic acid to reduce the methanol concentration to ≤5%. Blank samples, i.e. samples without starting material, were prepared the same way as the tissue samples.

Solid phase extraction was carried out on a Pressure+ 48 (positive pressure Manifold, Biotage) at ≤ 2 psi. SPE columns were pre-wet with 1 mL methanol and then equilibrated with 2 mL wash buffer [5% (v/v) methanol, 0.1% (v/v) acetic acid]. Samples spiked with internal standard mixtures of endocannabinoids (20 µL) and oxylipins (10 µL) were added to the columns. After washing with 2 mL wash buffer, the samples were eluted with 3 mL acetonitrile, followed by 2 ml methanol. Finally, the samples were dried using a vacuum concentrator (miVac Quattrol Concentrator, Genevac) and reconstituted in 110 μL of 2.27 μg/L 12-[[(cyclohexylamino)carbonyl]amino]-dodecanoic acid in methanol.

Liquid chromatography-mass spectrometry was conducted using an Agilent UHPLC system (Infinity 1290) coupled with an electrospray ionization source (ESI) to an Agilent 6490 triple quadrupole system equipped with iFunnel Technology (Agilent Technologies, Santa Clara, CA, USA). Metabolite separation was performed using a Waters BEH C18 column (2.1 mm × 150 mm, 130 Å, 1.7-μm particle size). The mobile phase consisted of (A) 0.1% acetic acid in milli-Q water and (B) acetonitrile:isopropanol (90:10). A flow rate of 300 μL/min and 10-μL injection volumes were employed for each run. The endocannabinoids are easily degraded compared to the relatively stable oxylipins wherefore all samples were first injected for ionization in positive mode (endocannabinoids) followed by all samples injected for negative (oxylipins) mode.

Quantification of the compounds was performed with MassHunter Quantitative Analysis QQQ (Agilent, Santa Clara, CA, USA).

### Statistical analyses

Results were analyzed with Prism 9.0.2 (GraphPad, San Diego, CA-US). The Kolmogorov-Smirnov test was used for normality testing. Then, results were either analyzed with the Mann-Whitney test to compare ranks, or analyzed using unpaired, 2-tailed Students t-test or ANOVA followed by independent comparisons with the Fisher’s least significant difference (LSD) test. Significance was accepted for P<0.05. Partial least-squares discriminant analysis (PLS-DA) was applied on z-scores of transcriptomics or lipidomics data sets using MATLAB 2019a (MathWorks, Natick, MA-USA). Results are presented as mean±SD unless otherwise stated. Statistical details of experiments can be found in the figure legends.

## Results

### HSL is enriched in neuronal synapses and is widely distributed in the brain

From the mouse cortex, we prepared synaptosomes, which are resealed nerve terminals after tissue homogenization. Synaptosomes were subsequently fractioned into synaptic preparations that are rich in PSD95 (post-synaptic density protein 95), SNAP25 (synaptosomal-associated protein 25) and synaptophysin, which correspond to the post-, pre- and extra-synaptic zones, respectively (figure 1A). Upon immunoblotting, we observed 2 bands at the approximate molecular weight of previously described HSL isoforms (Lindvall *et al*., 2004), but only the band corresponding to the predominant isoform in adipose tissue (84 kDa) was absent from in protein extracts from HSL-/- mice (figure 1B). The synaptic fractions showed a more intense immunoreactivity against HSL than total protein extracts (figure 1B). Indeed, synaptosomes showed 8-fold more immunoreactivity than total extracts (P<0.01) and, among the synaptic preparations, HSL immunoreactivity was observed in pre- and mostly post-synaptic fractions (figure 1C). We then set to investigate the cerebral distribution of HSL. First, we measured DAG lipase activity in the absence or presence of an anti-HSL antibody that abrogates HSL activity. Although HSL-specific DAG lipase activity was 20-fold lower in brain than in adipose tissue, is was detected in all brain areas analysed, and on average it accounted for about 2/3 of total DAG lipase activity (figure 1D). Then, the presence of HSL in these mouse brain samples was confirmed by immunoblotting experiments (figure 1E). Different unspecific bands in blots of figures 1B and 1E is attributed to difference in sample preparation.

**Figure 1.**
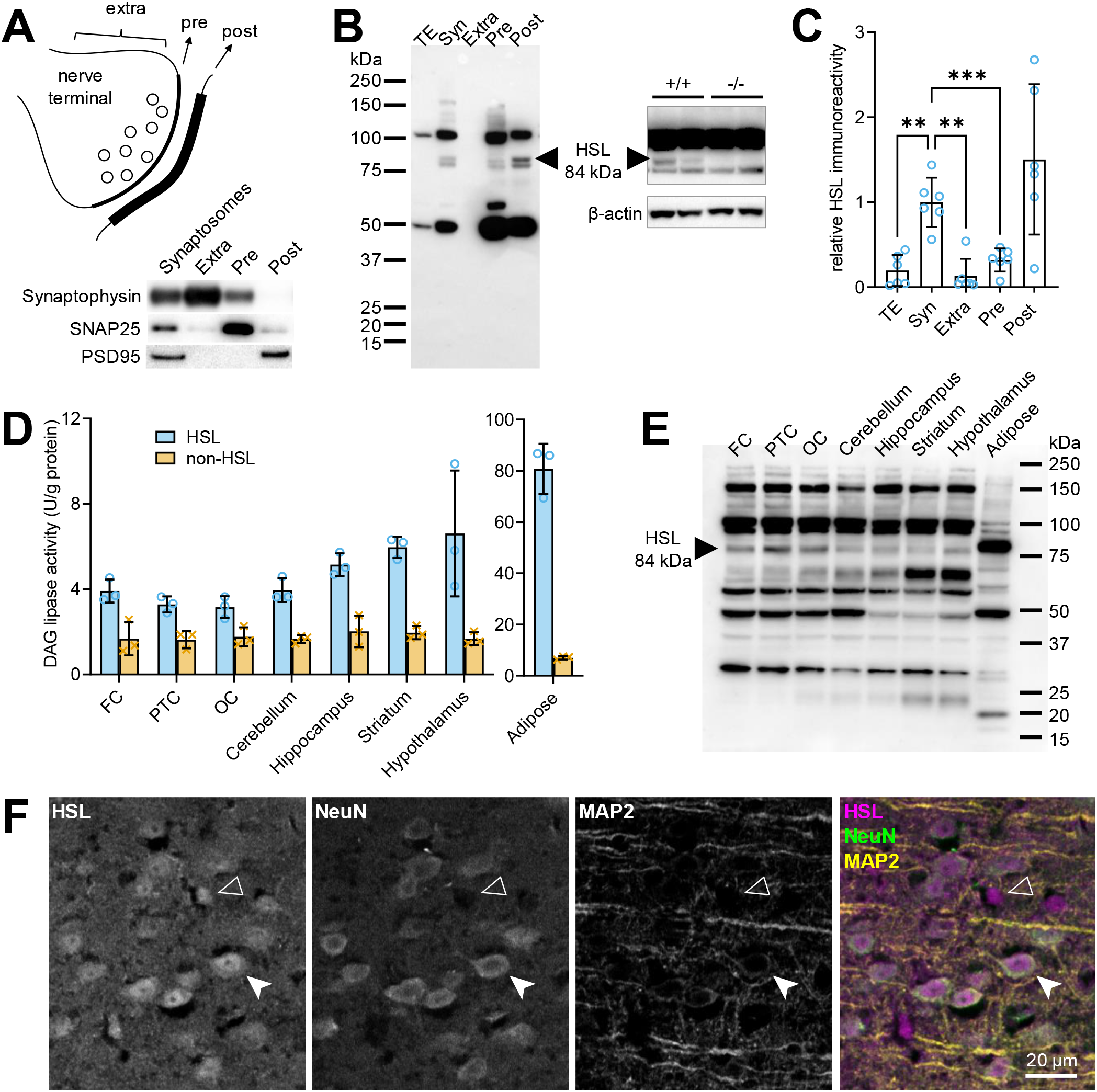
HSL is present in neurons and distributed throughout the mouse brain. (A) Western blots of pre-synaptic (Pre), post-synaptic (Post) and extra-synaptic (Extra) fractions prepared from the mouse cortex show enrichment in SNAP25, PSD95 and synaptophysin, respectively. (B) Relative to total protein extracts (TE), synaptosomes, pre- and post-synaptic fractions show immunoreactivity against HSL, a band of 84 kDa that is absent from protein extracts of the HSL-/- mouse HSL-/- mouse cortex. Protein loaded in gels was 30 µg for synaptic fractions and 40 µg for HSL-/- testing. (C) Quantitative analysis of western blot immunoreactivity suggests pre- and mostly post-synaptic HSL enrichment. (D-E) HSL-specific (blue circles) and non-specific (orange crosses) DAG lipase activity in brain areas and adipose tissue (D) and respective immunoreactivity in western blotting (E). Abbreviations for cortical areas: FC, frontal cortex; PTC, parieto-temporal cortex; OC, occipital cortex. Protein loaded for immunoblotting was 40 µg for brain samples, and 15 µg for adipose tissue. (F) Representative fluorescence micrographs of the mouse brain cortex immunolabeled for HSL (magenta), NeuN (green) and MAP2 (yellow). HSL immunoreactivity appears within NeuN-positive (filled arrowhead) and NeuN-negative (open arrowhead) cells. Data in bar graphs are represented as individual data points and mean±SD. Symbols over data-points indicate significant differences between control and HSL-/- mice (**P<0.01 and ***P<0.001) based on Fisher’s LSD post hoc comparison after ANOVA.

Immunofluorescence microscopy in mouse brain slices showed that HSL immunoreactivity is observed within all cells that are positive for the widely used neuronal marker NeuN, as well as in NeuN-negative cells (figure 1F), suggesting that both neurons and glia express HSL. In cultured cortical neurons, HSL immunoreactivity was observed in the cell soma and along dendritic processes (figure 2A). In finer dendrites, HSL immunoreactivity was adjacent to that of PSD95 (figure 2B), in line with its synaptic location.

**Figure 2.**
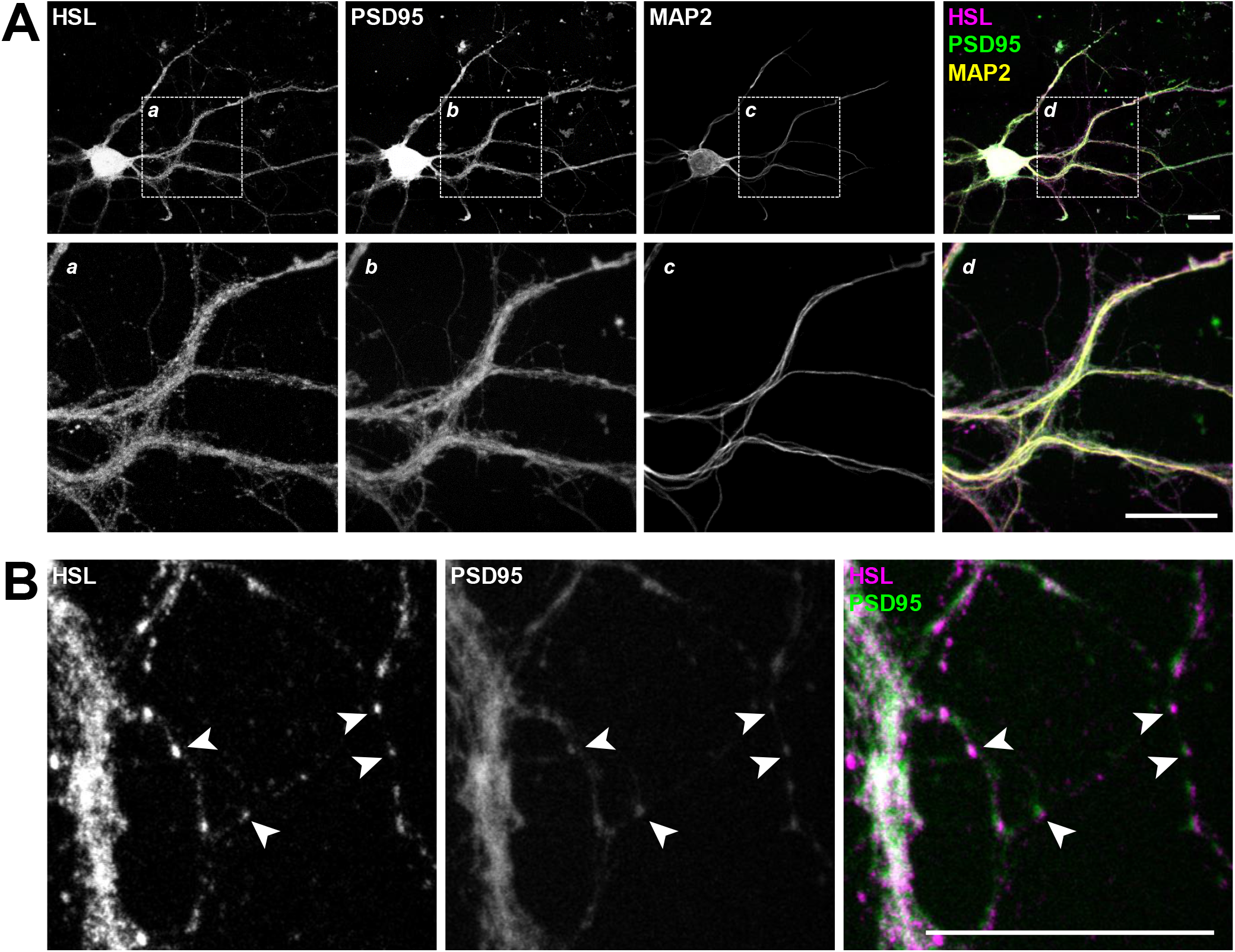
HSL is observed in primary cultured neurons. (A) Representative immunofluorescence micrographs of cultured cortical neurons stained for HSL (magenta), PSD95 (green) and MAP2 (yellow) showing HSL immunoreactivity throughout the cell soma and axons. (B) Higher magnification images show adjacent immunoreactivity against HSL and PSD95 in fine neuronal processes (arrowheads). Scale bars in micrographs are 20 µm.

### Characteristics of HSL -/- mice

Mice in this study were 19-22 months old, and HSL-/- mice showed slightly lower body weight (29.6±4.9 *versus* 36.7±6.6 g, P<0.05), higher fed plasma insulin (1874±1005 *versus* 701±782 pmol/L, P<0.01) but similar plasma glucose (10.0±1.3 *versus* 10.8±2.1 mmol/L, P<0.05) compared to the corresponding wild-type littermates.

### HSL null mice display short- and long-term memory impairment

To investigate whether HSL contributes to brain function, HSL null (-/-) mice and wild-type littermates (HSL+/+) were analysed using a battery of behaviour tests that probe for memory, exploratory behaviour and locomotor activity, and anxiety-like behaviour.

Spatial learning and memory was analysed using the Barnes maze, in which mice were trained for 8 days to learn the escape box location, that is the target hole, and long-term memory was analysed after 48 hours (figure 3A). Learning during the 8 training days was indistinct between genotypes (figure 3B). In training day 1, the experiment was preceded by a habituation session in which mice were placed in the escape whole, that is the target, and released in the center of the arena 2 hours later. Thereafter, the first training session revealed short-term memory impairment in HSL -/- mice, as depicted by larger latency (P<0.05) and number of errors until the target was reached (P<0.05), when compared to wild-type mice (figure 3C). During a training period of 8 days, HSL-/- mice utilized more often a random than serial hole search to identify the target (figure 3D). Long-term memory was probed 48 hours later, with mice being allowed to explore the maze without the escape box. HSL-/- mice searched less holes in the target quadrant (P<0.05), took longer time to reach the target hole for the first time (P<0.05), and spent overall less time in the target quadrant than controls (P<0.01, figure 3E), suggesting impaired spatial memory performance. The reduced Y-maze spontaneous alternation in HSL-/- mice compared to controls (P<0.05) further confirmed memory impairment caused by genetic HSL deletion (figure 3F).

**Figure 3.**
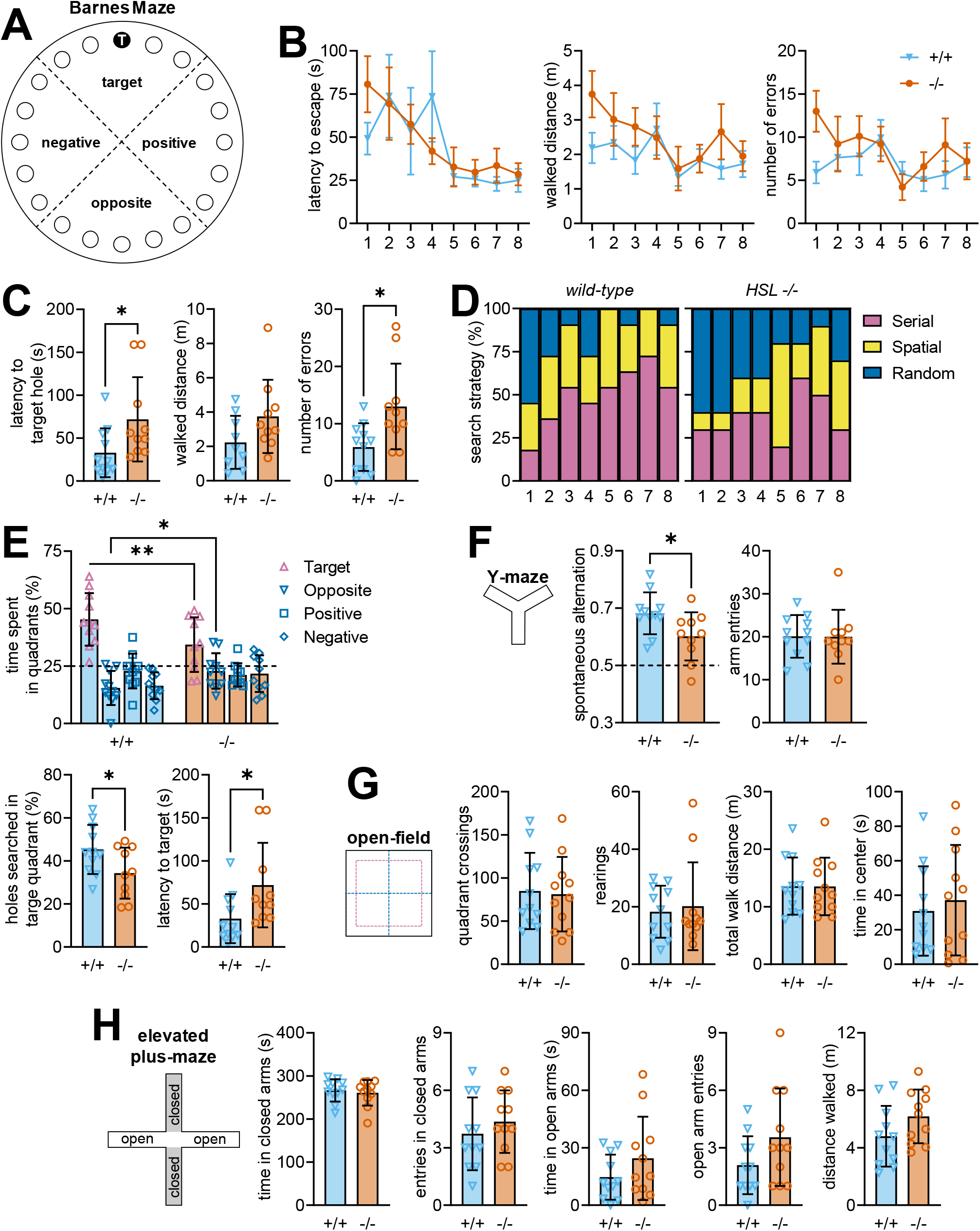
HSL -/- mice show memory impairment while exploratory and anxiety-like behaviours are preserved. (A) Schematic representation of the Barnes maze with holes numbered 1 to 10 from the target hole (T), and divided in 4 quadrants. (B) Learning to find the target hole over the 8 acquisition days was not affected by genotype. (C) Training in acquisition day 1 was followed after habituation to the maze, and allowed to identify increased latency and walked distance until entering the target hole, an increased number of errors (non-target hole searches) in HSL-/- *versus* wild-type mice. (D) Compared to controls, HSL-/- mice utilized more often a random than serial hole search during the 8-day training period. (E) Relative to controls, HSL-/- mice searched less holes in the target quadrant, took longer time to reach the target hole for the first time, and spent overall less time in the target quadrant during the probe session. Dashed line at 25% represents random exploration (chance). (F) Fraction of successful spontaneous alternations (consecutive entries in the 3 arms) in the Y-maze was lower in HSL-/- mice than controls, while showing similar number of entries in the maze arms. Dashed line at 0.5 represents random arm entries. (G) Genotype had no effect on open-field exploration, with similar number of quadrant crossings, rearing events (vertical explorations), walked distance in the arena, and time spent in the arena center. (H) Elevated plus-maze showed no effect of genotype on either time spent and number of entries in closed and open arms, or total distance walked in the maze. Data are represented as mean±SEM in line graphs of n=10-11, and as individual data points and mean±SD in bar graphs. Symbols over data-points indicate significant differences between control and HSL-/- mice (*P<0.05 and **P<0.01) based on Student t-tests or Fisher’s LSD post hoc comparison following presence of significant effects of genotype in ANOVA tests.

These memory assessments were not biased by alterations of exploratory behaviour since the genotype had no effect on the number of entries in the Y-maze arms (figure 3F), or on any measure of the open-field exploration, such as quadrant crossings, rearing and total walked distance in the arena (figure 3G). HSL deletion was also unrelated to development of anxiety-like behaviours depicted by changes in the exploration of the unprotected open-field arena center (figure 1G) or of the open arms in the elevated plus-maze (figure 3H).

### HSL deletion impacts genes involved in glucose metabolism and inflammation

As a first step to investigate the impact of HSL deletion on brain metabolism, we measured the expression of a panel of genes coding for proteins involved in glucose and lipid metabolism and genes encoding enzymes involved in the endocannabinoid system. Since neuroinflammation is known to be involved in a number of neurological and metabolic disorders (de Bem *et al*., 2021), this gene panel also included the master regulator of inflammatory processes NF-κB and cytokines. Transcriptomics data from the hippocampus and cortex was analysed with a 2-component partial least-squares discriminant analysis (PLS-DA), and allowed good separation of HSL-/- and wild-type mice (figure 4A), with PC1 and PC2 accounting for 72% and 16% of the variance in gene expression. The PLS-DA model coefficients (figure 4B) indicates that, in the hippocampus, HSL deletion increases the expression of genes necessary for glucose metabolism, namely the glucose carriers GLUT1, 3 and 4, as well as Peroxisome proliferator-activated receptor gamma (PPARγ) and Peroxisome proliferator-activated receptor gamma coactivator 1-α (PGC1α) that are key in mitochondria physiology. The hippocampus of HSL-/- mice also showed reduced expression of hexokinase (HK1) and a number of genes involved in lipid metabolism (DAGLα, ABDH6, MAGL, ATGL and LPL). Some of these observations were paralleled in the cortex. However, most strikingly, the cortex showed a prominent increase of NF-κB and the pro-inflammatory interleukins IL-6 and IL-1β (IL-1β was also increased in hippocampus), suggesting the occurrence of neuroinflammation.

**Figure 4.**
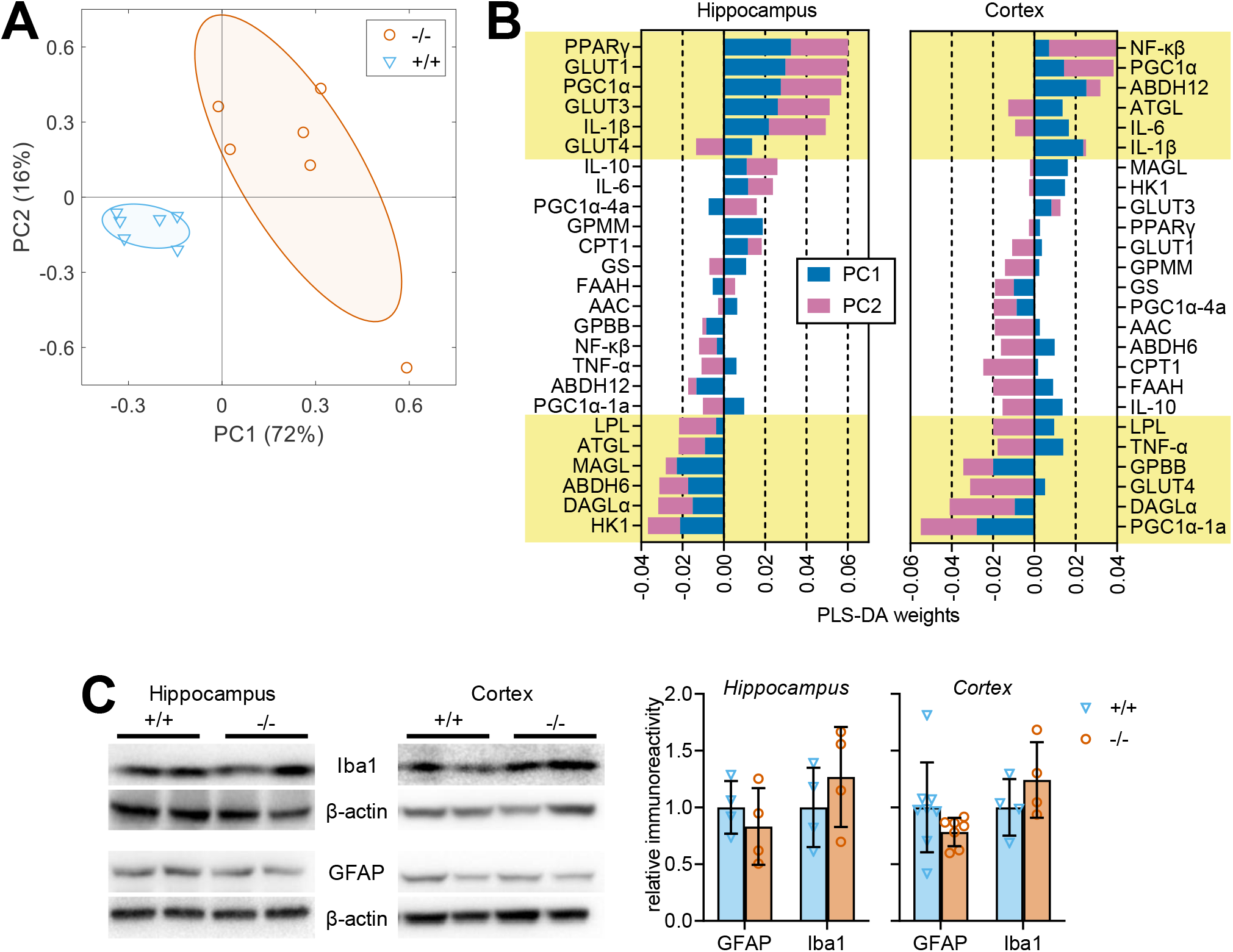
Effect of HSL deletion on cortical and hippocampal expression of genes involved in in inflammation and metabolism. (A) Transcriptomics on 12 mice revealed distinct gene expression between HSL+/+ and -/- mice in a PLS-DA, with PC1 and PC2 accounting for 72% and 16% of the total variance. Ellipsoids delineate SD. (B) The weights of PC1 and PC2 with most differentially expressed genes in the hippocampus and cortex highlighted in yellow. Genes are ordered by absolute sum of PC1 and PC2 weights, and taking in account the sign of the largest coefficient. (C) Relative immunoreactivity of the microglial marker Iba-1 and astrocytic marker GFAP in total protein extracts of the cortex and hippocampus (n=4-6). Immunoreactivity is normalised to the average of controls. Data are represented as individual data points and mean±SD in bar graphs.

Given the association of HSL deletion to neuroinflammation, we then tested whether this process is accompanied by gliosis. Immunoblotting against the astrocytic marker GFAP (glial fibrillary acidic protein) or the microglial marker Iba1 (allograft inflammatory factor 1) showed similar immunoreactivity in cortical or hippocampal extracts from HSL -/- and +/+ mice (figure 4C). Altogether, this suggests that HSL deletion is associated to low-grade neuroinflammation without the occurrence of important astrogliosis or microgliosis.

### HSL deletion does not impact typical markers of synaptic health

Since HSL is located in synapses, we then tested whether memory impairment upon HSL deletion was caused by loss of synaptic proteins involved in neurotransmitter release, as observed in various models of metabolic disease (Garcia-Serrano & Duarte, 2020). Protein extracts from the cortex showed reduced immunoreactivity against syntaxin-4 in HSL-/- mice compared to controls (P<0.05), without substantial effects on syntaxin-1, SNAP25 or synaptophysin (figure 5A). In the hippocampus the density of these four proteins was similar in the two genotypes (figure 5B).

**Figure 5.**
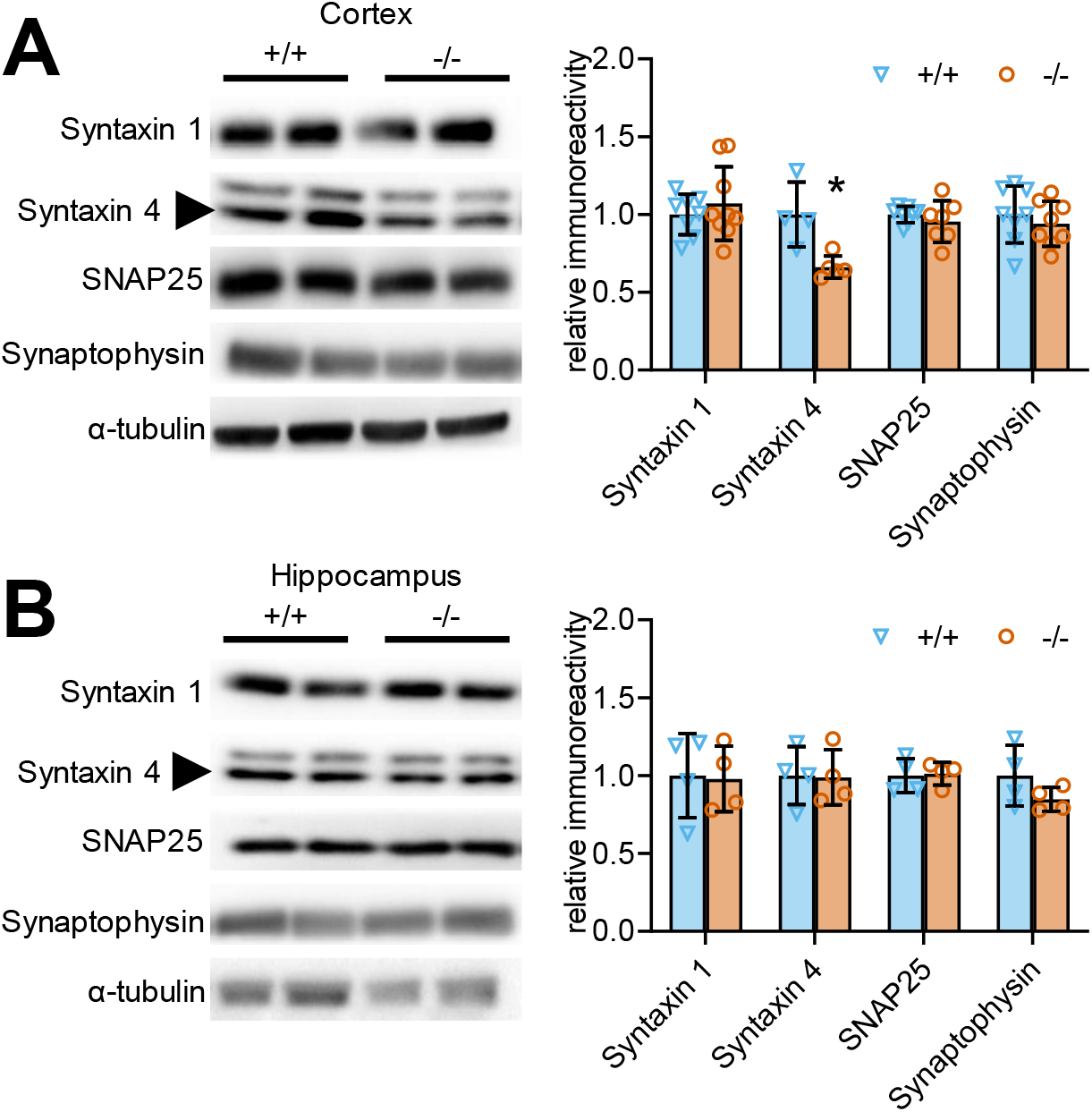
Effect of HSL deletion on protein markers of synaptic health. Relative immunoreactivity of syntaxin-1 (35 kDa), syntaxin-4 (34 kDa; arrow), SNAP25 (25 kDa) and synaptophysin (37 kDa), which are key proteins involved in neurotransmitter release, in total protein extracts of the cortex (A) and hippocampus (B). Immunoreactivity is normalised to the average of controls. Data are represented as individual data points and mean±SD of n=4-8 in bar graphs. Symbols over data-points indicate significant differences between control and HSL-/- mice (*P<0.05) based on 2-tailed Student t-tests.

### HSL deletion causes a shift in the profile of available bioactive lipids

Given the particular synaptic enrichment of HSL in synapses, it is plausible that cellular signalling through HSL is involved in the control of lipid metabolites that modulate synaptic physiology. Thus, in a further set of experiments, we determined the concentration of endocannabinoids and oxylipins in the hippocampus and cortex. A panel of over 50 bioactive lipids was analysed by mass spectrometry (tables 1-2), and we then applied a PLS-DA to determine the most important set of metabolites differentiating between HSL+/+ and -/- mice. By using 2 principal components (PC) we were able to observe a reasonable separation of the 2 genotypes (figure 6A), with PC1 and PC2 accounting for 32% and 36% of the variance in lipids. The PLS-DA model coefficients (figure 6B-C) allowed us to identify a number of bioactive lipids of importance in the genotype discrimination, including the endocannabinoids 2-linoleoyl glycerol (2-LG), *N*-arachidonoyl glycine (NAGly), arachidonyl etanolamide (AEA) and docosahexaenoyl ethanolamide (DHEA), the prostaglandins D2 (PGD2), E2 (PGE2) and F2α (PGF2a), and other eicosanoids such as (12S)-12-hydroxyeicosapentaenoic acid [12(S)-HEPE], 5,6-epoxyeicosatrienoic acid [5(6)-EpETrE], 11,12-epoxyeicosatrienoic acid [11(12)-EpETrE], 9-hydroxy-5Z,7E,11Z,14Z-eicosatetraenoic acid [(+-)9-HETE], 20-hydroxyarachidonic acid (20-HETE), leukotriene B4 (LTB4) and *trans-*leukotriene B4 (trans-LTB4), or thromboxane B2 (TxB2). When analysing concentrations of the lipids individually, however, no significant impact of HSL deletion was observed (figures 6D-E)

**Figure 6.**
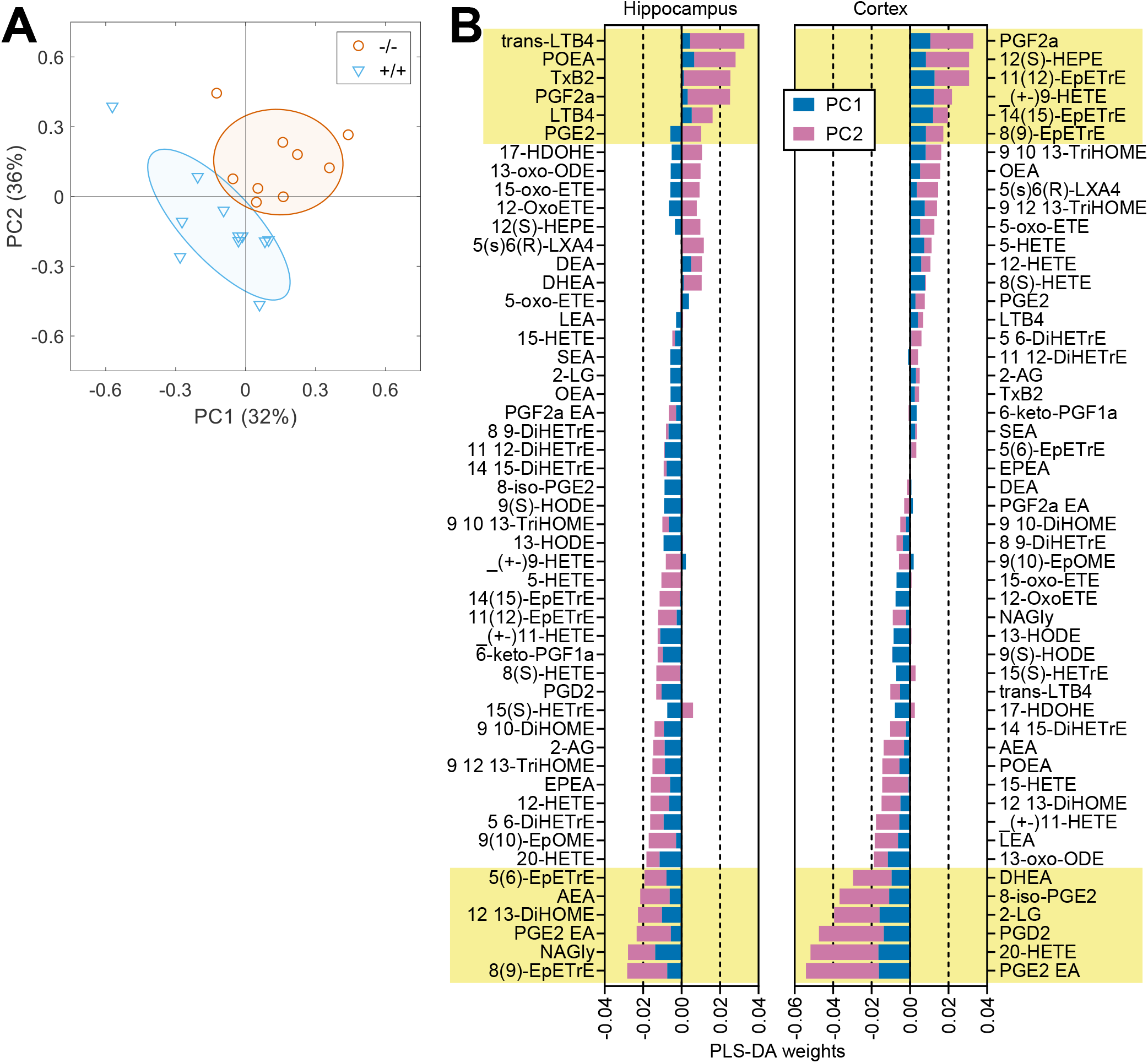
Effect of HSL deletion on bioactive lipids. (A) A panel of bioactive lipids analysed by mass spectrometry allowed to discriminate between HSL+/+ and -/- mice in a PLS-DA. PC1 and PC2 accounted for 32% and 36% of the total variance. Ellipsoids represent SD. (B) The weights of PC1 and PC2 identify a number of bioactive lipids of importance in the PLS-DA. Lipids in stacked bar graphs are ordered by absolute sum of PC1 and PC2 weights, and taking in account the sign of the largest coeficient. For complete metabolites abbreviations, see STAR methods. *Six most important hippocampal lipids with positive coefficients:* Trans-Leukotriene B4 (trans-LTB4), Palmitoleoyl ethanolamide (POEA), Thromboxane B2 (TxB2), Prostaglandin F2α (PGF2a), Leukotriene B4 (LTB4), Prostaglandin E2 (PGE2). *Six most important hippocampal lipids with negative coefficients:* 8,9-epoxyeicosatrienoic acid [8(9)-EpETrE], *N*-Arachidonoyl glycine (NAGly), Prostaglandin E2 ethanolamide (PGE2 EA), 12,13-Dihydroxyoctadec-9(Z)-enoic acid (12,13-DiHOME), Arachidonyl etanolamide (AEA), 5,6-epoxyeicosatrienoic acid [5(6)-EpETrE]. *Six most important cortical lipids with positive coefficients:* Prostaglandin F2α (PGF2a), (12S)-12-hydroxyeicosapentaenoic acid [12(S)-HEPE], 11,12-epoxyeicosatrienoic acid [11(12)-EpETrE], 9-hydroxy-5Z,7E,11Z,14Z-eicosatetraenoic acid [(+-)9-HETE], 14,15-epoxyeicosatrienoic acid [14(15)-EpETrE], 8,9-epoxyeicosatrienoic acid [8(9)-EpETrE]. *Six most important cortical lipids with negative coefficients:* Prostaglandin E2 ethanolamide (PGE2 EA), 20-Hydroxyarachidonic acid (20-HETE), Prostaglandin D2 (PGD2), 2-Linoleoyl glycerol (2-LG), 8-iso Prostaglandin E2 [8-iso-PGE2], Docosahexaenoyl ethanolamide (DHEA).

## Discussion

Our results indicate that HSL is active throughout the brain and accounts for at least two thirds of total DAG lipase activity throughout the assessed brain areas. Accordingly, HSL-/- mice were reported to have a substantial dampening of brain DAG lipase activity (Haemmerle *et al*., 2002). It should be noted that our HSL activity assay is optimised for measuring HSL. It might be that the relative DAG lipase activity accounted from HSL is lower in the native tissue, which allows for further on-demand activation. Similarly, Blankman *et al* (2007) have assayed lipase activity under conditions that are far from optimal for HSL, and concluded that HSL has negligible 2-AG hydrolysis activity. Therefore, further studies are necessary for the determination of the exact roles of brain HSL.

Furthermore, the present study suggests that HSL is particularly located in synapses, where it might exert important regulatory mechanisms that are hitherto undescribed. For the first time to our knowledge, we demonstrate here that HSL is necessary for memory performance, since the HSL-/- mouse exhibited impaired short- and long-term memory in the Barnes maze, as well as impaired spatial working memory in the Y-maze. This role of HSL in cognition adds up to previous suggestions of HSL-mediated lipolysis modulating hypothalamic function (Sekiya *et al*., 2004; Hundahl *et al*., 2021).

Given the synaptic enrichment of HSL, we evaluated whether degeneration of cortical and hippocampal synapses was a possible cause for memory impairment, as in other models of metabolic and neurological disorders (Martinsson *et al*., 2019; Garcia & Duarte, 2020). Levels of synaptophysin, syntaxin-1 or SNAP25, which are enriched in nerve terminals were not modified by HSL deletion. HSL deletion caused a reduction of syntaxin-4 levels in the cortex (but not hippocampus). Although this protein is part of the exocytosis machinery of nerve terminals, it is mainly present in processes of astrocytes (Tao-Cheng *et al*., 2015). Therefore, it is possible that release of molecules from astrocytes acting on synapses is impaired. Nevertheless, we have not observed alterations in the levels of GFAP that is specific to astrocytes. Neuronal communication at synapses is crucial for cognition, and damage to synapses and impaired synaptic functions is associated to cognitive decline (Morrison and Baxter, 2012; Sheng *et al*., 2012). Although HSL-/- mice do not show loss of synaptic markers that is typical of neurodegeneration, one cannot exclude impairment in the modulation of neurotransmitter release.

Neuroinflammation is a key process by which lipid metabolism disorders and obesity impact brain function (de Bem *et al*., 2021). HSL-/- mice show low grade neuroinflammation characterized by higher expression of pro-inflammatory cytokines than wild-type littermates. This occurs mainly in the hippocampus, and thus might contribute to the observed spatial memory impairment. Immunofluorescence experiments revealed HSL expression in NeuN-negative cells, which are likely astrocytes or microglia. Moreover, the concentration of oxylipins involved in inflammatory processes were modified upon genetic HSL deletion (see below). Low-grade inflammation in adipose tissue of HSL-/- mice has been described (Hansson et al, 2006). It has also been shown that HSL deficiency alters the expression of elongases and desaturases in liver and adipose tissue (Fernandez *et al*, 2011; Morigny *et al*, 2019), which might impact lipogenesis and the profile of circulating long-chain polyunsaturated fatty acids from which inflammatory mediators are synthesized. Therefore, given the pivotal participation of lipid metabolism in microglia modulation (Layé *et al*., 2018), future studies should further address a putative role of HSL in controlling neuroinflammatory cues.

The possibility of functional rather than structural derangement in synapses, as well as the low-grade inflammation, suggests that HSL is a key regulator of neuroactive lipid products. Indeed, we have identified overall basal changes in the concentration of eicosanoids and endocannabinoids induced by HSL deletion in the cortex and hippocampus. Most of these functional lipid products originate from arachidonic acid, which is a relatively abundant fatty acid in the brain (Contreras *et al*., 2000). In addition to being the primary hydrolytic enzyme for 2-AG, the most abundant endocannabinoid in the brain (Dinh *et al*, 2002), MAGL is considered to be the rate-limiting enzyme for release of arachidonic acid in the brain, to be used for the synthesis of neuroinflammatory prostaglandins and other eicosanoids, namely prostaglandin E2 (PGE2), PGD2, PGF2α and thromboxane B2 (TxB2) (Nomura *et al*, 2011). These contributed to the differences observed in the profile of bioactive lipids between HSL-/- and +/+ mice, suggesting that HSL might also have a role in the production of such eicosanoids.

Eicosanoids also participate in the modulation of cerebral blood flow in response to neuronal demands by acting on the vascular bed (Atwell *et al*., 2010; Sonnay *et al*., 2017). The lack of matching nutrient supply from the circulation to the demands of neuronal work is certainly critical for adequate functional performance. Cellular adaptations might be in place to cope with the lack of vascular flexibility, and our analysis found for example the increase in expression of the glucose carrier GLUT1, that is ubiquitous in blood vessels and brain cells, as well as the neuronal specific GLUT3, and the insulin-sensitive GLUT4 that is particularly located near synapses (Ashrafi *et al*., 2017).

Together, MAGL, ABHD12, and ABHD6 control about 99% of 2-AG signalling in the brain (Savinainen *et al*., 2012), and each enzyme exhibits a distinct subcellular distribution, suggesting that they regulate distinct pools of 2-AG in the nervous system (Blankman *et al*., 2007). Expression of ABHD12 and ABHD6 was modified in cortex and hippocampus, respectively. Furthermore, MAGL expression tended to be increased in the cortex and reduced in the hippocampus to nearly half of that in wild-type mice. Despite small, MAGL expression changes upon HSL deletion might still be critical in the dampening of endocannabinoid signalling in confined compartments such as the synapse, namely for removal of 2-AG produced following neurotransmitter release. Indeed, genetic deletion of MAGL results in the accumulation of 2-AG and other MAG species in the brain, without changes in expression of HSL, ABHD6 and ABHD12 (Taschler *et al*., 2011). None of the most studied endocannabinoids - 2-AG and anandamide - were modified in the hippocampus and cortex by HSL deletion. However, it should be noted that our lipid profile was measured in the resting brain, not in response to increased neuronal activity that is the trigger for endocannabinoid synthesis and release (Metna-Laurent & Marsicano, 2015).

FAAH hydrolyzes not only anandamide but other bioactive lipids, such as *N*-palmitoylethanolamine (POEA) (Fezza *et al*., 2008), which does not bind cannabinoid receptors, but functions as an endogenous ligand for peroxisome proliferator receptor-α (PPAR-α) (Lo Verme *et al*., 2005). Although no FAAH expression changes were induced by HSL deletion, POEA concentration was larger in the hippocampus of HSL-/- than +/+ mice. POEA acting on PPAR-α might play a role in mitochondrial physiology and, indeed, we observed brain alterations of PGC1-α or its isoforms in HSL-/- mice.

DAGL-α deletion in mice was also proposed to disrupt learning and memory due to the depletion of 2-AG and arachidonic acid across the whole brain (Schurman *et al*., 2019). Interestingly, Schurman *et al*. found effects of DAGL-α deletion on brain anandamide levels that were not mimicked by pharmacological inhibition of the enzyme, indicating that enzyme activities producing and/or degrading anandamide are also impacted by deletion of DAGL-α. In our study, we measured expression of such enzymes, and found negligible effects triggered by HSL deletion.

In sum, we found no signs of substantial synaptic degeneration, astrogliosis or microgliosis, but there was a mild increase in levels of pro-inflammatory cytokines. The profile of endocannabinoids and eicosanoids was distinct in the hippocampus and cortex of HSL-/- and +/+ mice, which might underlie the observed memory impairment. Therefore, we conclude that HSL is a key modulator of eicosanoids that are immuno-modulators, and further propose a role for HSL on the neuro- and vaso-active lipid synthesis upon demand by increased neuronal activity. The later remains to be demonstrated.

## Abbreviations

2-AG: 2-arachidonoylglycerol
ATGL: adipose triglyceride lipase
DAG: diacylglycerol
FAAH: fatty acid amide hydrolase
HSL: hormone-sensitive lipase
IL: interleukin
MAGL: monoacylglycerol lipase
PC: principal components
PG: prostaglandin
RT-PCR: real-time polymerase chain reaction
TAG: triacylglycerol

## Acknowledgement

We thank Dr. Isak Martinsson and Prof. Gunnar K. Gouras for providing the cell cultures for this study and Sara Larsson for technical assistance with the HSL activity measurements. Lund University Bioimaging Centre is acknowledged for providing access to microscopy resources. Swedish Metabolomics Centre (Umeå, Sweden) is acknowledged for the measurement of oxylipins and endocannabinoids.

This study was conducted with funding from the Tage Blücher Foundation, Dementiafonden, Swedish foundation for International Cooperation in Research and Higher education (BR2019-8508), Swedish Research council (2019-01130), Diabetesfonden (Dia2019-440), Crafoord Foundation (20190007, 20200564), and Direktör Albert Påhlssons Foundation (FB2018-0265, FB2019-0321). J.D. acknowledges generous financial support from The Knut and Alice Wallenberg foundation, the Faculty of Medicine at Lund University and Region Skåne. The authors acknowledge support from the Lund University Diabetes Centre, which is funded by the Swedish Research Council (Strategic Research Area EXODIAB, grant 2009-1039) and the Swedish Foundation for Strategic Research (grant IRC15-0067).

## Author Contributions

C.H. and J.D. designed the study. C.H., C.S. and J.D. performed experiments, analyzed data, and wrote and revised the manuscript.

